# Searching the *Pinus taeda* foliar mycobiome for emerging pathogens among brown spot needle blight and needlecast outbreaks in the Southeast United States

**DOI:** 10.1101/2024.03.13.584907

**Authors:** Colton D. Meinecke, Afaq M.M. Niyas, Elizabeth McCarty, Tania Quesada, Jason A. Smith, Caterina Villari

## Abstract

Needle pathogens cause the discoloration, death, or premature abscission of conifer foliage, which reduce growth and vigor, and repeated defoliation may eventually result in tree mortality. Since 2016, forest managers in the southeast United States have reported an increasing scale, frequency, and severity of needle disease outbreaks on the region’s principal timber species, loblolly pine (*Pinus taeda* L.). These recent outbreaks are raising concern throughout the region, as needle diseases are not traditionally considered a threat to *P. taeda*. *Lecanosticta acicola* (Thum.) Syd., the native causal agent of brown-spot needle blight, has been recovered from some outbreaks, however, the full array of fungi associated with symptoms has not been explored. In this research, *P. taeda* foliage was collected from affected stands throughout the region and analyzed to identify fungi associated with needle disease symptoms. We employed both targeted molecular diagnostics, to confirm the presence or absence of *L. acicola*, and DNA metabarcoding, to characterize the foliar mycobiome and screen for other potential pathogens. *Lecanosticta acicola* was detected among symptomatic needles from multiple states, particularly in western portions of the *P. taeda* range but rarely from stands in eastern states. Fungal ITS1 metabarcoding revealed multiple pathogens in symptomatic needles and identified associations between known pathogens fungi and differing symptoms. Additionally, the fungal community composition of needles varied with patterns of symptom presentation. This study is the first regionwide assessment of fungi associated with recent large-scale needle disease outbreaks on *P. taeda* and identifies multiple pathogens that warrant study in greater detail.

## Introduction

Healthy southern pine forests dominate southeastern landscapes and are critical to the ecology and economy of the Southeast United States (Oswalt et al. 2019). Southern pines are commonly planted and managed for productivity in working forests, maintaining native forest cover while offering employment for local communities and provide forest products, revenue, and a means to sequester carbon (Greene et al. 2019; Zhao et al. 2016). Loblolly pine (*Pinus taeda* L.) is the primary southern pine species in the region (Allen et al. 2005; Oswalt et al. 2019). An estimated 22 billion *P. taeda* grow throughout its range, comprising eight percent of total above-ground biomass in the United States (Oswalt et al. 2019). *Pinus taeda* is also the most widely planted tree in North America (Oswalt et al., 2019), with over 14 million hectares planted in the southeast United States (South & Harper, 2016). However, these forests now face a growing threat from a historically unimportant enemy: needle disease (Pandit et al. 2020).

The incursion, emergence, and damage caused by foliar pathogens in coniferous forests is on the rise worldwide (Bradshaw 2004; Broders et al. 2014; Costanza et al. 2018; Ganley et al. 2014; Hu et al. 2020; Theron et al. 2022; Tubby et al. 2023; van der Nest et al. 2019b). Foliar diseases of pines, referred to as needle blights or needlecasts, are caused by fungi and oomycetes that attack and kill photosynthetic tissue. The resulting defoliation impairs light capture, evapotranspiration, nutrient uptake, stem growth, and stand productivity, and contributes to tree mortality (Barnard and Ash 1998; Bednářová et al. 2013; Fraser et al. 2016; Jansons et al. 2020; Shaw et al. 2021). In disrupting host leaf function, the development of foliar disease also alters the mycobiome, or whole fungal community, of leaf tissues, exacerbating stress and promoting attack by opportunistic facultative pathogens (Ata et al. 2022; Busby et al. 2016; Ginnan et al. 2020; Ridout and Newcombe 2015).

Fungi known to attack *P. taeda* foliage include species in the genera *Hypoderma, Lophodermium, Rhizosphaera,* and *Lecanosticta* (Barnard and Ash 1998; Boyce 1954, 1969; Cunningham 2020; van der Nest et al. 2019a). Of note, *Lecanosticta acicola* (Thum.) Syd. is the well-described causal agent of brown-spot needle blight (BSNB) on *Pinus* spp. In North America, *L. acicola* is often observed causing BSNB on the grass stage of its coevolved host longleaf pine (*P. palustris* Mill.) (Siggers 1932; van der Nest *et al*. 2019b), and on introduced Austrian (*P. mugo* Turro) (Cleary et al. 2019) and Scots pines (*P. sylvestris* L.) (Skilling and Nicholls 1974). *Lecanosticta acicola* is also an associate of the recently observed white pine needle damage (WPND) on eastern white pine (*P. strobus* L.) (Broders *et al*. 2014). Damaging outbreaks of needle disease have also affected planted *P. taeda* in the past (Cordell et al. 1990; Edgerton and Moreland 1924), but such events were infrequent and considered unusual. Broadly speaking, foliar pathogens were considered weak opportunistic attackers on *P. taeda*, and the conditions they cause were rarely managed (Barnard and Ash 1998). However, reports from industry and government forest managers indicate a stark increase in needle disease frequency and severity, calling attention to an emerging threat for which the causal agents are poorly understood (Datta et al. 2021; Pandit et al. 2020).

Starting in 2016, forest health practitioners recorded increased needle disease damage on *P. taeda* in many southeastern states (Pandit *et al*. 2020). Recently reported needle disease symptoms include reddening or browning of needles from the bottom of the crown upward, premature needle drop, and mortality in the most severe cases. The needles often display irregular, gregarious necrotic spots or bands, which are surrounded by yellow chlorotic zones. Fungal fruiting bodies may be found erupting from within diseased tissue. Often, the necrotic zones merge and result in the death of the entire distal portion of needles. In some cases, needles appear necrotic without defined spots or bands. Foliar and whole-tree symptoms of needle diseases are visually similar, and the observed symptoms are consistent with those caused by many fungi (Barnard and Ash 1998). Large-scale needle disease outbreaks have been assessed among commercially managed *P. taeda* in the southeast United States, and *L. acicola* was detected from symptomatic needles from outbreaks in Alabama, Arkansas, Louisiana, and Mississippi (Datta et al. 2021; Meinecke et al. 2021). While *L. acicola* has been associated with some outbreaks, it has not been consistently recorded in all of the major pine needle disease outbreaks. Moreover, the extent of damage on *P. taeda* is unusual and is not typical of *L. acicola* infection on this host. Additionally, the observed symptoms may be caused by multiple separate diseases at the various outbreak locations, or even the result of a complex of fungi as is suggested in white pine needle defoliation in the Northeast US (Munck et al. 2011; Wyka et al. 2017).

A better characterization of the pathogens involved in this needle disease phenomenon is critical for understanding this emerging threat, developing management strategies, and ultimately preserving native southeastern pine forests. Advances in next-generation sequencing (NGS) have opened the door to exploring and characterizing mycobiomes, or entire fungal communities, and identifying potential pathogens in novel disease outbreaks (Bulman et al. 2018; Jongman et al. 2020; Newberry et al. 2023). DNA metabarcoding, or amplicon sequencing, is a high throughput sequencing technique that amplifies and sequences specific, conserved DNA regions from an environmental sample. Organisms within a sample can be sequenced simultaneously and analyzed as a community. Metabarcoding was largely a tool for ecological investigations, but it is increasingly used for surveillance and diagnosis of plant and animal disease due to its ability to reveal pathogens without the need of isolation or targeted molecular tools (Banchi et al. 2018; Huggins et al. 2019).

In this study, molecular diagnostics and next-generation sequencing were employed to address the emerging threat of needle disease on *P. taeda*. We outlined four hypotheses as follows: 1) The brown-spot needle blight pathogen *L. acicola* will be present among diseased *P. taeda* needles at outbreak sites. 2) The presence of certain fungal taxa will be correlated with needle disease symptoms. 3) The overall fungal communities within *P. taeda* foliage from affected stands will differ in composition (i.e., membership) and richness based on geographic location. 4) The fungal communities will be distinct among pines with different needle disease symptom presentations, differing in composition and taxa richness. To this aim, we analyzed needles from impacted stands throughout the southeast United States. First, we employed a loop-mediated isothermal amplification (LAMP)-based species-specific molecular assay (Aglietti et al. 2021) to determine the presence or absence of *L. acicola*. Then, using DNA metabarcoding, we comprehensively analyzed the fungal communities associated with needle samples and identified potential pathogens.

## Materials and Methods

### Sampling criteria

Sampling covered *P. taeda* forests across the southeast United States, including sites in Arkansas, Alabama, Florida, Georgia, Louisiana, Mississippi, South Carolina, and Texas (Table 1), where partners in state, federal, and private industrial organizations collected samples on our behalf. Sites were selected where outbreaks of visible foliar disease symptoms, including browning and premature drop of needles, were occurring on *P. taeda*. Sampling locations were not predetermined, rather a call to submit samples was sent to a broad network of partners. Samples to be processed were selected among those received to represent locations throughout the southeast United States and as funding permitted. Sampling date, geographic location, and symptom presentation were recorded for each sample at the time of collection. We received 195 samples from 98 trees across all sampling locations, and 161 of those samples from 92 trees could be analyzed.

**Table 1:** Samples analyzed for possible needle pathogens.

| Sample ID | Symptom Presentation Type<sup>a</sup> | State | County | Collection Date | LAMP | Metabarcoding |
| --- | --- | --- | --- | --- | --- | --- |
| 20210607-AL-A01 | Asymptomatic | Alabama | Crenshaw | 6/7/2021 | - | - |
| 20210607-AL-A02 | Asymptomatic | Alabama | Crenshaw | 6/7/2021 | - | + |
| 20210607-AL-A03 | Asymptomatic | Alabama | Crenshaw | 6/7/2021 | - | - |
| 20210607-AL-A04 | Asymptomatic | Alabama | Crenshaw | 6/7/2021 | - | - |
| 20210607-AL-A05 | Asymptomatic | Alabama | Crenshaw | 6/7/2021 | - | - |
| 20210607-AL-S01 | BSNB | Alabama | Crenshaw | 6/7/2021 | + | + |
| 20210607-AL-S02 | BSNB | Alabama | Crenshaw | 6/7/2021 | + | + |
| 20210607-AL-S03 | BSNB | Alabama | Crenshaw | 6/7/2021 | + | + |
| 20210607-AL-S04 | BSNB | Alabama | Crenshaw | 6/7/2021 | - | + |
| 20210607-AL-S05 | BSNB | Alabama | Crenshaw | 6/7/2021 | + | + |
| 20200701-AR-A01 | Asymptomatic | Arkansas | Saline | 7/1/2020 | - | + |
| 20200701-AR-A05 | Asymptomatic | Arkansas | Saline | 7/1/2020 | - | + |
| 20200701-AR-S01 | BSNB | Arkansas | Saline | 7/1/2020 | + | + |
| 20200701-AR-S05 | BSNB | Arkansas | Saline | 7/1/2020 | + | + |
| 20200710-AR-A02 | Asymptomatic | Arkansas | Garland | 7/10/2020 | - | + |
| 20200710-AR-A04 | Asymptomatic | Arkansas | Garland | 7/10/2020 | - | + |
| 20200710-AR-A05 | Asymptomatic | Arkansas | Garland | 7/10/2020 | + | + |
| 20200710-AR-S02 | BSNB | Arkansas | Garland | 7/10/2020 | - | + |
| 20200710-AR-S04 | BSNB | Arkansas | Garland | 7/10/2020 | + | + |
| 20200710-AR-S05 | BSNB | Arkansas | Garland | 7/10/2020 | + | + |
| 20210121-AR-A03 | Asymptomatic | Arkansas | Saline | 1/21/2021 | - | - |
| 20210121-AR-A04 | Asymptomatic | Arkansas | Saline | 1/21/2021 | - | Dropped |
| 20210121-AR-A05 | Asymptomatic | Arkansas | Saline | 1/21/2021 | - | Dropped |
| 20210121-AR-S01 | BSNB | Arkansas | Saline | 1/21/2021 | + | + |
| 20210121-AR-S02 | BSNB | Arkansas | Saline | 1/21/2021 | + | + |
| 20210121-AR-S03 | BSNB | Arkansas | Saline | 1/21/2021 | - | Dropped |
| 20210121-AR-S04 | BSNB | Arkansas | Saline | 1/21/2021 | - | Dropped |
| 20210121-AR-S05 | BSNB | Arkansas | Saline | 1/21/2021 | - | Dropped |
| 20210304-AR-A01 | Asymptomatic | Arkansas | Howard | 3/4/2021 | - | + |
| 20210304-AR-A02 | Asymptomatic | Arkansas | Howard | 3/4/2021 | - | - |
| 20210304-AR-A04 | Asymptomatic | Arkansas | Howard | 3/4/2021 | - | + |
| 20210304-AR-A05 | Asymptomatic | Arkansas | Howard | 3/4/2021 | - | - |
| 20210304-AR-S01 | BSNB | Arkansas | Howard | 3/4/2021 | - | + |
| 20210304-AR-S02 | BSNB | Arkansas | Howard | 3/4/2021 | - | + |
| 20210304-AR-S03 | BSNB | Arkansas | Howard | 3/4/2021 | - | - |
| 20210304-AR-S04 | BSNB | Arkansas | Howard | 3/4/2021 | - | Dropped |
| 20210304-AR-S05 | BSNB | Arkansas | Howard | 3/4/2021 | - | - |
| 20210311-AR-A01 | Asymptomatic | Arkansas | Montgomery | 3/11/2021 | - | + |
| 20210311-AR-A03 | Asymptomatic | Arkansas | Montgomery | 3/11/2021 | - | + |
| 20210311-AR-A05 | Asymptomatic | Arkansas | Montgomery | 3/11/2021 | - | - |
| 20210311-AR-S01 | BSNB | Arkansas | Montgomery | 3/11/2021 | - | Dropped |
| 20210311-AR-S02 | BSNB | Arkansas | Montgomery | 3/11/2021 | + | + |
| 20210311-AR-S03 | BSNB | Arkansas | Montgomery | 3/11/2021 | - | - |
| 20210311-AR-S04 | BSNB | Arkansas | Montgomery | 3/11/2021 | - | + |
| 20210311-AR-S05 | BSNB | Arkansas | Montgomery | 3/11/2021 | - | Dropped |
| 20210322-AR-A01 | Asymptomatic | Arkansas | Howard | 3/22/2021 | - | Dropped |
| 20210322-AR-A02 | Asymptomatic | Arkansas | Howard | 3/22/2021 | - | + |
| 20210322-AR-A03 | Asymptomatic | Arkansas | Howard | 3/22/2021 | - | + |
| 20210322-AR-A04 | Asymptomatic | Arkansas | Howard | 3/22/2021 | - | - |
| 20210322-AR-A05 | Asymptomatic | Arkansas | Howard | 3/22/2021 | - | + |
| 20210322-AR-S01 | BSNB | Arkansas | Howard | 3/22/2021 | + | + |
| 20210322-AR-S02 | BSNB | Arkansas | Howard | 3/22/2021 | - | + |
| 20210322-AR-S03 | BSNB | Arkansas | Howard | 3/22/2021 | + | Dropped |
| 20210322-AR-S04 | BSNB | Arkansas | Howard | 3/22/2021 | - | + |
| 20210322-AR-S05 | BSNB | Arkansas | Howard | 3/22/2021 | - | + |
| 20210624-AR-A01 | Asymptomatic | Arkansas | Grant | 6/24/2021 | - | Dropped |
| 20210624-AR-A02 | Asymptomatic | Arkansas | Grant | 6/24/2021 | - | - |
| 20210624-AR-A03 | Asymptomatic | Arkansas | Grant | 6/24/2021 | - | - |
| 20210624-AR-A04 | Asymptomatic | Arkansas | Grant | 6/24/2021 | - | - |
| 20210624-AR-A05 | Asymptomatic | Arkansas | Grant | 6/24/2021 | - | Dropped |
| 20210624-AR-S01 | BSNB | Arkansas | Grant | 6/24/2021 | - | - |
| 20210624-AR-S02 | BSNB | Arkansas | Grant | 6/24/2021 | - | - |
| 20210624-AR-S03 | BSNB | Arkansas | Grant | 6/24/2021 | + | + |
| 20210624-AR-S04 | BSNB | Arkansas | Grant | 6/24/2021 | - | - |
| 20210624-AR-S05 | BSNB | Arkansas | Grant | 6/24/2021 | - | - |
| 20191205-FL-A01 | Asymptomatic | Florida | Alachua | 12/5/2019 | - | - |
| 20191205-FL-A02 | Asymptomatic | Florida | Alachua | 12/5/2019 | - | - |
| 20191205-FL-S01 | Needlecast | Florida | Alachua | 12/5/2019 | - | - |
| 20191205-FL-S02 | Needlecast | Florida | Alachua | 12/5/2019 | - | - |
| 20200702-FL-A21 | Asymptomatic | Florida | Alachua | 7/2/2020 | - | - |
| 20200702-FL-A22 | Asymptomatic | Florida | Alachua | 7/2/2020 | - | - |
| 20200702-FL-A23 | Asymptomatic | Florida | Alachua | 7/2/2020 | - | - |
| 20200702-FL-A24 | Asymptomatic | Florida | Alachua | 7/2/2020 | - | - |
| 20200702-FL-A25 | Asymptomatic | Florida | Alachua | 7/2/2020 | - | - |
| 20200702-FL-A26 | Asymptomatic | Florida | Alachua | 7/2/2020 | - | - |
| 20200702-FL-S21 | Needlecast | Florida | Alachua | 7/2/2020 | - | - |
| 20200702-FL-S22 | Needlecast | Florida | Alachua | 7/2/2020 | - | - |
| 20200702-FL-S23 | Needlecast | Florida | Alachua | 7/2/2020 | - | - |
| 20200702-FL-S24 | Needlecast | Florida | Alachua | 7/2/2020 | - | - |
| 20200702-FL-S25 | Needlecast | Florida | Alachua | 7/2/2020 | - | - |
| 20200702-FL-S26 | Needlecast | Florida | Alachua | 7/2/2020 | - | - |
| 20210507-GA-A01 | Asymptomatic | Georgia | Appling | 5/7/2021 | - | - |
| 20210507-GA-A02 | Asymptomatic | Georgia | Appling | 5/7/2021 | - | Dropped |
| 20210507-GA-A03 | Asymptomatic | Georgia | Appling | 5/7/2021 | - | - |
| 20210507-GA-A04 | Asymptomatic | Georgia | Appling | 5/7/2021 | - | - |
| 20210507-GA-A05 | Asymptomatic | Georgia | Appling | 5/7/2021 | - | - |
| 20210507-GA-S01 | Needlecast | Georgia | Appling | 5/7/2021 | - | - |
| 20210507-GA-S02 | Needlecast | Georgia | Appling | 5/7/2021 | - | - |
| 20210507-GA-S03 | Needlecast | Georgia | Appling | 5/7/2021 | - | - |
| 20210507-GA-S04 | Needlecast | Georgia | Appling | 5/7/2021 | - | - |
| 20210507-GA-S05 | Needlecast | Georgia | Appling | 5/7/2021 | - | - |
| 20220315-GA-A01 | Asymptomatic | Georgia | Hart | 3/15/2022 | - | - |
| 20220315-GA-A02 | Asymptomatic | Georgia | Hart | 3/15/2022 | - | - |
| 20220315-GA-A03 | Asymptomatic | Georgia | Hart | 3/15/2022 | - | - |
| 20220315-GA-A04 | Asymptomatic | Georgia | Hart | 3/15/2022 | - | - |
| 20220315-GA-A05 | Asymptomatic | Georgia | Hart | 3/15/2022 | - | + |
| 20220315-GA-S01 | Needlecast | Georgia | Hart | 3/15/2022 | - | - |
| 20220315-GA-S02 | Needlecast | Georgia | Hart | 3/15/2022 | - | - |
| 20220315-GA-S03 | Needlecast | Georgia | Hart | 3/15/2022 | - | - |
| 20220315-GA-S04 | Needlecast | Georgia | Hart | 3/15/2022 | - | - |
| 20220315-GA-S05 | Needlecast | Georgia | Hart | 3/15/2022 | - | - |
| 20220519-GA-A01 | Asymptomatic | Georgia | Gilmer | 5/19/2022 | - | + |
| 20220519-GA-A02 | Asymptomatic | Georgia | Gilmer | 5/19/2022 | - | Dropped |
| 20220519-GA-A03 | Asymptomatic | Georgia | Gilmer | 5/19/2022 | - | - |
| 20220519-GA-A04 | Asymptomatic | Georgia | Gilmer | 5/19/2022 | - | - |
| 20220519-GA-S01 | BSNB | Georgia | Gilmer | 5/19/2022 | - | + |
| 20220519-GA-S02 | BSNB | Georgia | Gilmer | 5/19/2022 | - | + |
| 20220519-GA-S03 | BSNB | Georgia | Gilmer | 5/19/2022 | - | - |
| 20220519-GA-S04 | BSNB | Georgia | Gilmer | 5/19/2022 | - | - |
| 20220610-LA-A01 | Asymptomatic | Louisiana | Grant Parish | 6/10/2022 | + | + |
| 20220610-LA-A02 | Asymptomatic | Louisiana | Grant Parish | 6/10/2022 | + | + |
| 20220610-LA-A03 | Asymptomatic | Louisiana | Grant Parish | 6/10/2022 | + | + |
| 20220610-LA-A04 | Asymptomatic | Louisiana | Grant Parish | 6/10/2022 | + | + |
| 20220610-LA-A05 | Asymptomatic | Louisiana | Grant Parish | 6/10/2022 | - | - |
| 20220610-LA-S01 | BSNB | Louisiana | Grant Parish | 6/10/2022 | + | + |
| 20220610-LA-S02 | BSNB | Louisiana | Grant Parish | 6/10/2022 | + | + |
| 20220610-LA-S03 | BSNB | Louisiana | Grant Parish | 6/10/2022 | + | + |
| 20220610-LA-S04 | BSNB | Louisiana | Grant Parish | 6/10/2022 | + | + |
| 20220610-LA-S05 | BSNB | Louisiana | Grant Parish | 6/10/2022 | + | + |
| 20221103-MS-A01 | Asymptomatic | Mississippi | Choctaw | 11/3/2022 | - | Dropped |
| 20221103-MS-A02 | Asymptomatic | Mississippi | Choctaw | 11/3/2022 | - | + |
| 20221103-MS-A03 | Asymptomatic | Mississippi | Choctaw | 11/3/2022 | - | + |
| 20221103-MS-A04 | Asymptomatic | Mississippi | Choctaw | 11/3/2022 | - | Dropped |
| 20221103-MS-A05 | Asymptomatic | Mississippi | Choctaw | 11/3/2022 | - | + |
| 20221103-MS-S01 | BSNB | Mississippi | Choctaw | 11/3/2022 | - | - |
| 20221103-MS-S02 | BSNB | Mississippi | Choctaw | 11/3/2022 | - | + |
| 20221103-MS-S03 | BSNB | Mississippi | Choctaw | 11/3/2022 | - | + |
| 20221103-MS-S04 | BSNB | Mississippi | Choctaw | 11/3/2022 | - | - |
| 20221103-MS-S05 | BSNB | Mississippi | Choctaw | 11/3/2022 | - | - |
| 20210429-SC-A01 | Asymptomatic | South Carolina | Saluda | 4/29/2021 | - | - |
| 20210429-SC-A02 | Asymptomatic | South Carolina | Saluda | 4/29/2021 | - | - |
| 20210429-SC-A03 | Asymptomatic | South Carolina | Saluda | 4/29/2021 | - | + |
| 20210429-SC-A04 | Asymptomatic | South Carolina | Saluda | 4/29/2021 | - | - |
| 20210429-SC-A05 | Asymptomatic | South Carolina | Saluda | 4/29/2021 | - | - |
| 20210429-SC-S01 | Needlecast | South Carolina | Saluda | 4/29/2021 | - | - |
| 20210429-SC-S02 | Needlecast | South Carolina | Saluda | 4/29/2021 | - | - |
| 20210429-SC-S03 | Needlecast | South Carolina | Saluda | 4/29/2021 | - | - |
| 20210429-SC-S04 | Needlecast | South Carolina | Saluda | 4/29/2021 | - | - |
| 20210429-SC-S05 | Needlecast | South Carolina | Saluda | 4/29/2021 | - | - |
| 20210819-SC-A01 | Asymptomatic | South Carolina | Greenwood | 8/19/2021 | - | - |
| 20210819-SC-A02 | Asymptomatic | South Carolina | Greenwood | 8/19/2021 | - | - |
| 20210819-SC-A03 | Asymptomatic | South Carolina | Greenwood | 8/19/2021 | - | - |
| 20210819-SC-A04 | Asymptomatic | South Carolina | Greenwood | 8/19/2021 | - | - |
| 20210819-SC-A05 | Asymptomatic | South Carolina | Greenwood | 8/19/2021 | - | Dropped |
| 20210819-SC-S01 | Needlecast | South Carolina | Greenwood | 8/19/2021 | - | - |
| 20210819-SC-S02 | Needlecast | South Carolina | Greenwood | 8/19/2021 | - | - |
| 20210819-SC-S03 | Needlecast | South Carolina | Greenwood | 8/19/2021 | - | - |
| 20210819-SC-S04 | Needlecast | South Carolina | Greenwood | 8/19/2021 | - | - |
| 20210819-SC-S05 | Needlecast | South Carolina | Greenwood | 8/19/2021 | - | - |
| 20210820-SC-A01 | Asymptomatic | South Carolina | Hampton | 8/20/2021 | - | - |
| 20210820-SC-A02 | Asymptomatic | South Carolina | Hampton | 8/20/2021 | - | - |
| 20210820-SC-A03 | Asymptomatic | South Carolina | Hampton | 8/20/2021 | - | - |
| 20210820-SC-A04 | Asymptomatic | South Carolina | Hampton | 8/20/2021 | - | - |
| 20210820-SC-A05 | Asymptomatic | South Carolina | Hampton | 8/20/2021 | - | - |
| 20210820-SC-S01 | Needlecast | South Carolina | Hampton | 8/20/2021 | - | + |
| 20210820-SC-S02 | Needlecast | South Carolina | Hampton | 8/20/2021 | - | - |
| 20210820-SC-S03 | Needlecast | South Carolina | Hampton | 8/20/2021 | - | - |
| 20210820-SC-S04 | Needlecast | South Carolina | Hampton | 8/20/2021 | - | - |
| 20210820-SC-S05 | Needlecast | South Carolina | Hampton | 8/20/2021 | - | - |
| 20220609-TX-A01 | Asymptomatic | Texas | Hardin | 6/9/2022 | - | + |
| 20220609-TX-A02 | Asymptomatic | Texas | Hardin | 6/9/2022 | - | - |
| 20220609-TX-A03 | Asymptomatic | Texas | Hardin | 6/9/2022 | - | + |
| 20220609-TX-A04 | Asymptomatic | Texas | Hardin | 6/9/2022 | - | + |
| 20220609-TX-A05 | Asymptomatic | Texas | Hardin | 6/9/2022 | - | + |
| 20220609-TX-S01 | BSNB | Texas | Hardin | 6/9/2022 | + | + |
| 20220609-TX-S02 | BSNB | Texas | Hardin | 6/9/2022 | - | - |
| 20220609-TX-S03 | BSNB | Texas | Hardin | 6/9/2022 | + | + |
| 20220609-TX-S04 | BSNB | Texas | Hardin | 6/9/2022 | + | + |
| 20220609-TX-S05 | BSNB | Texas | Hardin | 6/9/2022 | - | + |
| 20220629-TX-A01 | Asymptomatic | Texas | Polk | 6/29/2022 | - | - |
| 20220629-TX-A02 | Asymptomatic | Texas | Polk | 6/29/2022 | - | Dropped |
| 20220629-TX-A03 | Asymptomatic | Texas | Polk | 6/29/2022 | - | - |
| 20220629-TX-A04 | Asymptomatic | Texas | Polk | 6/29/2022 | - | - |
| 20220629-TX-A05 | Asymptomatic | Texas | Polk | 6/29/2022 | - | Dropped |
| 20220629-TX-A06 | Asymptomatic | Texas | Polk | 6/29/2022 | - | - |
| 20220629-TX-S01 | Needlecast | Texas | Polk | 6/29/2022 | - | - |
| 20220629-TX-S02 | Needlecast | Texas | Polk | 6/29/2022 | - | - |
| 20220629-TX-S03 | Needlecast | Texas | Polk | 6/29/2022 | - | - |
| 20220629-TX-S04 | Needlecast | Texas | Polk | 6/29/2022 | - | - |
| 20220629-TX-S05 | Needlecast | Texas | Polk | 6/29/2022 | - | - |
| 20220629-TX-S06 | Needlecast | Texas | Polk | 6/29/2022 | - | - |

<sup>a</sup> Symptom type was classified as brown spot needle blight (BSNB) symptoms, needlecast symptoms, or asymptomatic when they did not display any disease symptoms.
<sup>b</sup> Detection of *Lecanosticta acicola* using loop-mediated isothermal amplification was determined by amplification of pathogen DNA from pine needle total DNA extracts in at least two of three replicate reactions using a species-specific LAMP assay per Aglietti et al. (2021).
<sup>c</sup> Detection of *Lecanosticta acicola* by metabarcoding was determined by the presence of ITS1 sequence reads from the sample that matched to the pathogen.

A minimum of five trees were sampled at each site and both symptomatic samples and asymptomatic samples were collected from each tree. Each sample consisted of 10 or more individual needles. Pine foliage was considered symptomatic of needle disease if bands or mottled spots of discoloration, necrotic lesions, death of distal portions of the needle, or fungal fruiting bodies in diseased tissue were present. After collection, needle samples were individually bagged and shipped overnight under ambient conditions to the Forest Pathology Laboratory at the University of Georgia.

### Symptom classification and DNA extraction

Upon receipt, needles were immediately assessed, and the symptoms of each sample were classified based on overall presentation as follows: “Asymptomatic” needles were green and healthy, without evidence of necrosis or fungal reproductive structures. “BSNB” needles displayed gregarious brown necrotic spots or bands surrounded by yellow chlorotic zones, with occasional dead distal tips and dispersed black fungal acervuli, which are typical symptoms and signs of BSNB as caused by *L. acicola.* “Needlecast” samples showed distal tip death or necrosis of the entire needle with dispersed fruiting bodies of known fungi other than *L. acicola* and lacked necrotic spots (Figure 1).

**Figure 1:**
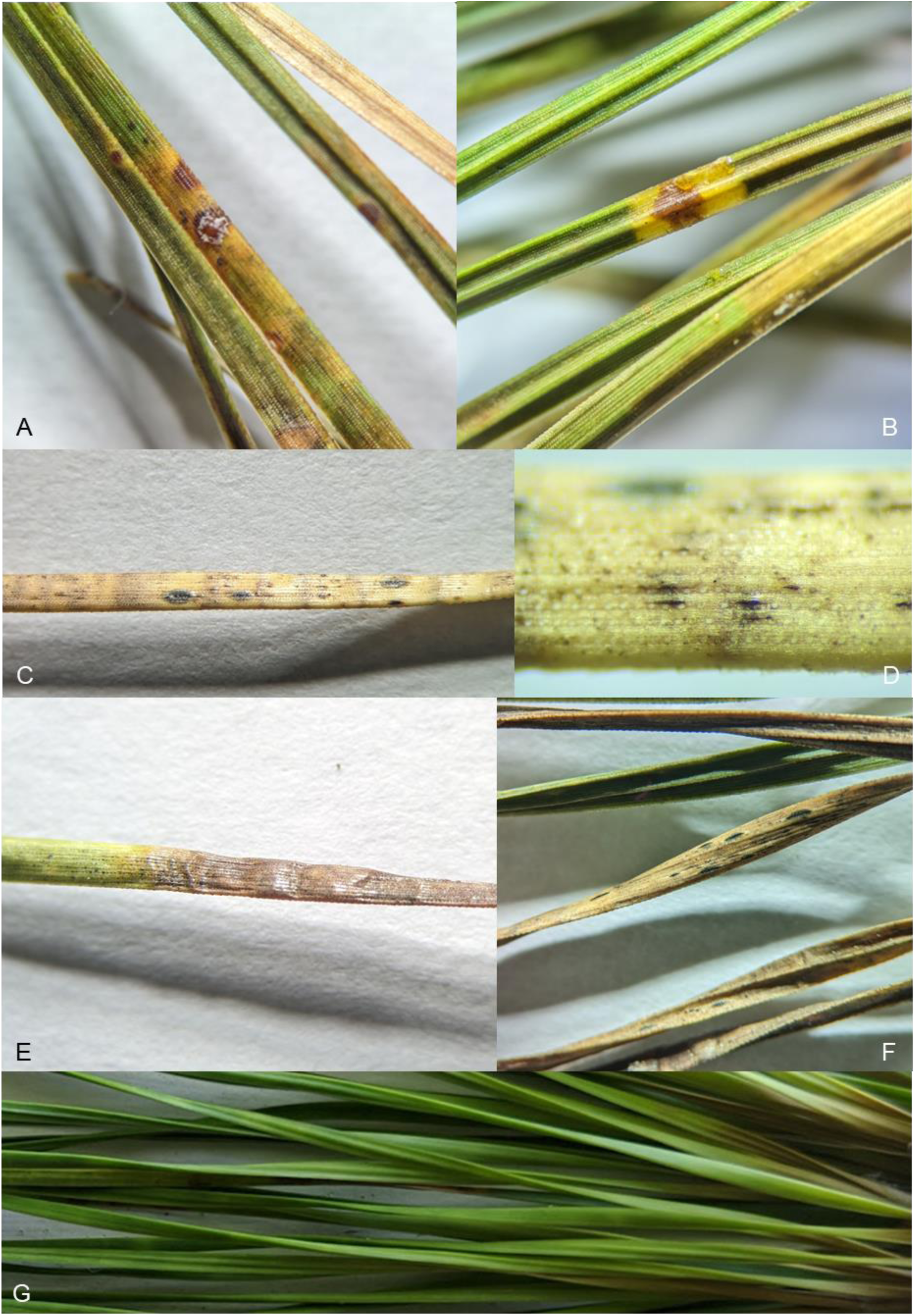
Examples of needle disease symptom presentation on *Pinus taeda* foliage. Samples were classified as displaying signs and symptoms of brown-spot needle blight, caused by *Lecanosticta acicola*, if they presented necrotic spots or bands surrounded in yellow chlorotic zones (A and B), with occasional dead distal tips (C and D) and fungal fruiting that included *L. acicola* acervuli (D). Samples were otherwise classified as needlecast if they displayed chlorosis, distal tip death or necrosis of the entire needle (E), and dispersed fungal fruiting, and which lacked *L. acicola* acervuli (F). Samples devoid of any sign or symptom of disease were classified as asymptomatic (G).

From each sample, ten needles were randomly selected for analysis. Each needle was surface sterilized by immersing in 95% ethanol (DeconLabs, King of Prussia, Pennsylvania, USA) for 5 seconds, 0.5% sodium hypochlorite (Clorox, Oakland, California, USA) for 2 minutes, and lastly 70% ethanol for 2 minutes (Oono et al. 2015). Needles were then air dried under sterile conditions and stored at -20°C prior to further processing. Each sample of ten needles was pooled and ground to a fine powder in a sterilized mortar and pestle with liquid nitrogen. Total DNA was extracted from 50.0 mg of fresh ground tissue using the DNeasy^®^ Plant Pro^®^ Kit (Qiagen, Germantown, Maryland, USA), following manufacturer instructions, and concentrations of DNA extracts were quantified using a Qubit fluorometer (Invitrogen, Carlsbad, California, USA). DNA extracts were stored frozen at -20 °C until analysis.

### Screening pine needles for the presence of L. acicola

Total DNA extracts were screened for the presence of *L. acicola* to rapidly diagnose cases of BSNB using a species-specific loop-mediated isothermal amplification (LAMP) assay (Aglietti et al. 2021). LAMP reactions utilized a FAM-labelled fluorescent reporter and were performed using a StepOnePlus^TM^ Real-Time PCR System (Applied Biosystems, Foster City, CA, USA). Reactions were assembled and conducted in triplicate as described by Aglietti et al. (2021). All LAMP primers, the FAM-labelled assimilating probe, and quencher strand were synthesized by Integrated DNA Technologies (IDT, Coralville, Iowa, USA). Each run included two non-template controls, in which molecular grade water was used in the place of template DNA, and DNA from *P. taeda* needles confirmed to be infected with *L. acicola* as a positive control. Samples were identified as infected by *L. acicola* if at least two out of three replicate LAMP reactions amplified successfully.

### Fungal isolates for metabarcoding mock communities

Four mock communities were included in each metabarcoding sequencing run, to serve as positive sequencing controls and to monitor for cross contamination during sample processing. Seven fungal isolates were used in total, and each mock community consisted of a different combination of the DNA from five of the seven fungi, such that each community was distinct from the others. DNA from each fungus of the mock community was present in equimolar amounts. The fungal isolates and the composition of each mock community are listed in Table 2. Fungal isolates were chosen because they were host-specific pathogens not known to inhabit pine tissues. It is critical to assemble mock communities that are composed of organisms unlikely to occur in test samples so that the occurrence of “index bleeding,” or reads incorrectly assigned to the wrong samples, can be quantified and bioinformatically mitigated (Tedersoo et al. 2022). Each fungal isolate was grown on sterile 2.0% potato dextrose agar (PDA, Becton Dickinson, Sparks, Maryland, USA) under ambient conditions for 14 days. From each isolate, approximately 100 mg of mycelia were collected, and total DNA was extracted using the E.Z.N.A.® Fungal DNA Mini Kit (Omega Bio-tek, Norcross, Georgia, USA) according to manufacturer instructions. Concentrations of DNA extracts were quantified using a Qubit fluorometer and normalized to 1.0 ng μL^−1^ prior to inclusion in mock communities.

**Table 2:** Isolates in each of the four mock communities included in the DNA metabarcoding sequence runs. DNA from each fungus was present in equimolar amounts. Mock communities served as positive sequencing controls and to monitor for cross contamination during each metabarcoding sequencing run.

|  | Mock 1 | Mock 2 | Mock 3 | Mock 4 |
| --- | --- | --- | --- | --- |
| Isolate 1 | *Harringtonia lauricola* | *Harringtonia lauricola* | *Harringtonia lauricola* | *Cryphonectria parasitica* |
| Isolate 2 | *Cryphonectria parasitica* | *Cryphonectria parasitica* | *Neofusicoccum carygenum* | *Neofusicoccum carygenum* |
| Isolate 3 | *Neofusicoccum carygenum* | *Neofusicoccum carygenum* | *Stagonosporopsis curcubitacearum* | *Stagonosporopsis curcubitacearum* |
| Isolate 4 | *Stagonosporopsis curcubitacearum* | *Exobasidium maculosum* | *Exobasidium maculosum* | *Exobasidium maculosum* |
| Isolate 5 | *Corynospora casiicola* | *Athelia rolfsii* | *Corynospora casiicola* | *Athelia rolfsii* |

### Fungal ITS1 metabarcoding and sequence analysis

The fungal communities of each needle sample were analyzed using DNA metabarcoding. The fungal rDNA internal transcribed spacer (ITS) genetic region is considered a universal barcode for identifying fungi in environmental samples and is the best-supported region for use in fungal diversity studies (Nilsson et al. 2019; Schoch et al. 2012). To enable discrimination between *Lecanosticta* species (van der Nest et al. 2019a), we targeted the ITS1 subregion using the primers ITS1f and ITS2, which effectively reduce the amplification of plant and bacterial DNA and are commonly utilized for fungal barcoding (Aguayo et al. 2018; Gardes and Bruns 1993; Schoch et al. 2012; White et al. 1990). In a preliminary test, we also attempted to additionally amplify the ITS2 subregion using the primers 5.8S-Fun and ITS4-Fun (Taylor et al. 2016). However, the primers amplified host plant (i.e., *P. taeda*) DNA such that plant sequences comprised most of the reads (data not shown), and this approach was abandoned. Total needle DNA extracts were submitted to the Georgia Genomics and Bioinformatics Core facility (GGBC, Athens, Georgia, USA) for library preparation and metabarcode sequencing on the Illumina® MiSeq platform (Illumina, San Diego, California, USA). Each of the three 96–well plate sequencing runs contained four mock communities, prepared as described above, and at least three blank DNA extracts to serve as positive and negative controls, respectively. Paired-end reads of 301 bp were generated using the MiSeq Reagent Kit v3 chemistry (600-cycle, Illumina).

Raw sequence reads were demultiplexed and converted to FASTQ format at the GGBC facility using Cassava v1.8 (Illumina). Following demultiplexing, all subsequent sequence processing was performed using the software package Quantitative Insights Into Microbial Ecology 2 v2023.2.0 (QIIME 2) in Python v3.8.8 (Bolyen et al. 2019; Van Rossum and Drake 2009), according to the QIIME2 Fungal ITS analysis guidelines (Bolyen et al. 2019; Huggins et al. 2019). In brief, the paired-end reads were merged, sequencing errors were corrected, reads were trimmed for quality with a threshold of Q20, and sequences that were short, low-quality, or chimeric, were discarded using the plugin DADA2 (Antich et al. 2021; Callahan et al. 2016; Huggins et al. 2019). Unique DNA sequences, termed exact sequence variants (ESVs), were identified by DADA2 (Callahan et al. 2017) and were taxonomically classified using a Naïve Bayes classifier, which is a machine learning model, trained on the UNITE fungal ITS dynamic reference database v8 (release 10.05.2021) (Nilsson et al. 2018). Count tables contained each ESV identified in every sample and the number of reads that matched that sequence. The counts were then randomly subsampled to 1676 counts per sample, to ensure even sampling and enable comparison (Weiss et al. 2017). This count number was chosen because it captured the highest number of ESVs while preserving most of the samples. Mock community sequences were processed separately following the same procedure. Reads from mock community samples were assessed to confirm successful sequencing and to gauge whether the proportion of reads reflected the proportions of DNA from fungi in each community.

### Statistical analyses

All data processing and statistical analyses were conducted in R, using R Studio (R Core Development Team, 2012). Low-frequency ESVs, defined as those with fewer than 10 observations across all samples, were pruned from the dataset. Sample count tables were then transformed to reflect the presence or absence of each fungus across all samples rather than attempting to quantify the abundance of each fungus. Deriving the abundance of a given species from fungal ITS1 is difficult due to the unknown amount of fungal biomass in the samples, differences in ITS copy number and length among species, and PCR bias (Tedersoo *et al*. 2022). The ESV counts, taxonomy data, and sample metadata were merged for analysis using *phyloseq* (McMurdie and Holmes 2013). The fungi identified from sample sequences were assessed to identify potential needle pathogens.

Multiple indicator species analyses (ISAs), one for each ESV, were performed on count table data to determine which individual fungal taxa correlate with symptom presentation (Dufrene and Legendre 1997; Morrison et al. 2021; Severns and Sykes 2020). Indicator values (IndVal) were calculated to measure the correlation between taxon presence and needle disease symptom presentation. Significant p-values were adjusted using the Benjamini-Hochberg method to control the false positivity rate in all indicator analyses (Benjamini and Hochberg 1995; Tedersoo et al. 2022; Yashiro et al. 2019). Because all asymptomatic samples were collected from symptomatic trees, and many pine needle pathogens are known to remain asymptomatic during latent infection periods, we excluded asymptomatic samples from the ISAs. Indicator species analyses were performed with 9,999 permutations in R using the *indicspecies::multipatt* function (De Cáceres et al. 2010).

The relationships of both geographic location and needle disease symptom presentation with fungal community composition and taxa richness response variables were tested with a permutational multivariate analysis of variance (PERMANOVA) and by comparing ESV richness, respectively. A Jaccard transformation was applied to binary count data to generate a matrix of dissimilarities between samples. The PERMANOVAs were performed on the dissimilarity matrix with 9,999 permutations (α = 0.05) using the *vegan::adonis* function (Oksanen et al. 2016), to determine whether fungal community composition, measured as the specific set of fungal ESVs in each sample, differed between types of symptom presentation or geographic location, and testing for interaction between these variables (McMurdie and Holmes 2013; Morrison et al. 2021; Skelton et al. 2019; Tedersoo et al. 2022).

The taxa richness of each sample was estimated using the observed number of ESVs, or exact sequence variants in each sample, a simple measure of the number of unique sequences recorded per sample as an additional means to compare fungal communities. The observed richness values were checked for normality using a Shapiro-Wilk test (Shapiro and Wilk 1965), and the means were compared between geographic location and needle disease symptom types using Kruskal-Wallis rank sum tests. Tests were followed by pairwise comparisons using a Wilcoxon rank sum test if a significant difference (α = 0.05) was detected (Ginnan *et al*. 2020).

## Results

### Symptom classification

We processed 195 needle samples collected from 98 *P. taeda* in 8 southeast states (Table 1). The number of trees sampled in each state varied, ranging from 30 from Arkansas to 5 each from Louisiana and Mississippi. In limited cases, only symptomatic foliage was present on the sampled trees at the time of collection, and no asymptomatic needles were collected in those instances. Symptomatic samples from Alabama, Arkansas, Louisiana, Mississippi and single sites in Georgia and Texas matched closely to the typical symptoms of BSNB and were classified accordingly, whereas symptomatic samples from the sites located in Florida, South Carolina, and the remaining sites in Georgia and Texas were classified as needlecast.

### Screening needles for L. acicola presence at outbreak sites using LAMP

The species-specific LAMP test detected *L. acicola* only from *P. taeda* that displayed BSNB symptoms (Table 1). *Lecanosticta acicola* positive samples included 10 of 30 trees sampled in Arkansas, 4 of 5 trees from Alabama, 4 of 5 from Louisiana, and 3 of 5 from the single BSNB site in Texas. In one case, both the symptomatic and asymptomatic samples from a single tree in Arkansas tested positive for *L. acicola* (Figure 2). Trees from Georgia, Mississippi, South Carolina, Florida, or the needlecast site in Texas were not positive for *L. acicola*.

**Figure 2:**
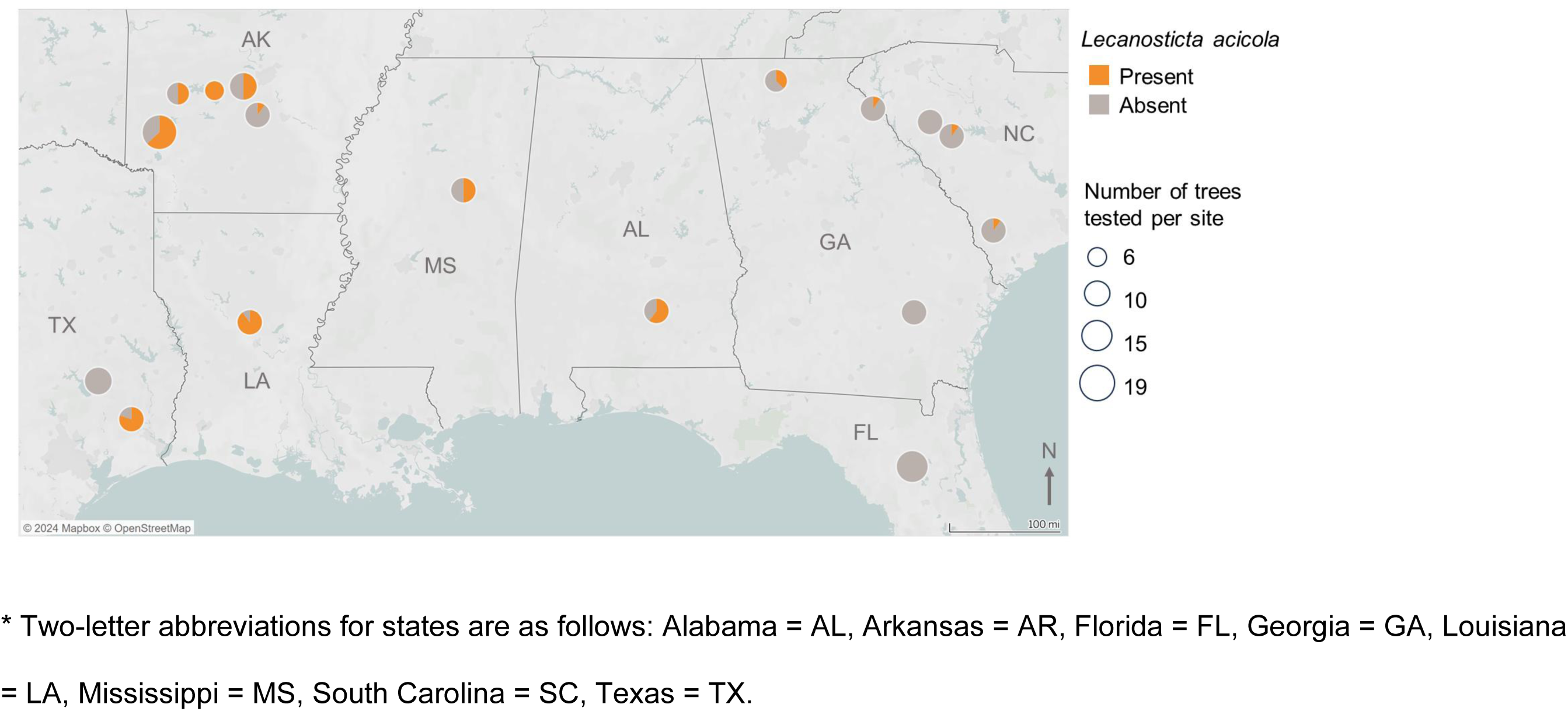
Sampling locations* at which *Pinus taeda* displayed symptoms of needle disease and the proportion of trees infected by *Lecanosticta acicola,* the causal agent of brown spot needle blight. Infection by *Lecanosticta acicola* was determined by positive amplification of pathogen DNA using a species-specific loop-mediated isothermal amplification assay or by metabarcoding through the presence of ITS1 sequence reads from the sample that matched the pathogen. The map was generated using Tableau version 2023.2 (Seattle, Washington, United States) from an OpenStreetMap (Mapbox, Cambridge, United Kingdom) template.

### Metabarcoding data quality

Illumina amplicon sequencing generated 27,213,358 (median 132,757) raw reads of the ITS1 subregion from needle DNA samples (Table 3). Denoising, filtering and merging reduced the ITS1 reads to 6,490,614 (median 13,540) and 12 samples were discarded due to low quality. Random subsampling to 1676 reads caused the removal of an additional 14 samples, which had too few reads to be analyzed. The final dataset contained evenly sampled reads, representing 169 samples from 97 trees. A total of 2,890 ESVs were identified, from which 176 taxa were classified.

**Table 3:** Number of fungal ITS1 sequence reads for all samples following each data processing step. Raw DNA sequences were filtered to remove primer and adapter sequences, and denoised to trim low-quality sequences and correct basecalling errors. Paired reads were merged and any reads too short to merge were discarded. Chimeric sequences, composed of erroneously combined fragments of separate sequences, were detected and discarded.

|  | Total raw | Filtered | Denoised<sup>a</sup> | Merged | Non-chimeric |
| --- | --- | --- | --- | --- | --- |
| Sum | 15,025,661 | 6,098,179 | 6,021,000 | 5,726,728 | 5,483,668 |
| Mean | 124,197.02 | 50,398 | 49,760.33 | 47,328.33 | 45,319.57 |
| Median | 90,459 | 29,765 | 28,461 | 27,679 | 24,484 |

<sup>a</sup> Denoising, paired-end sequence merging, and chimeric sequence detection were performed using DADA2 (Callahan *et al.* 2016).

Assessment of mock community reads confirmed heavy biases in the proportional representation of the included taxa, indicating that transformation to binary presence/absence data was necessary in this experiment. Sequences belonging to the mock community fungi could not be detected among needle sample data after quality control and random subsampling, indicating that read bleeding was minimal in the final dataset. Negative control blank samples were filtered out from the dataset during DADA2 processing due to insufficient high-quality reads, consistent with the absence of template DNA in these samples.

### Fungal taxa associated with P. taeda affected by needle disease

The identities of potential pathogens detected in pine needle samples are reported in Table 4. While the taxonomic classifier was trained using the curated UNITE database and produced high-confidence assignments, the identification of fungal taxa using a single locus can often lack resolution and the reported identities should all be considered preliminary (Ivanová et al. 2016). Several known needle pathogens were identified among sequences from both symptomatic and asymptomatic needles from sampled trees. Some known weak pathogens were identified in multiple locations, across different symptom classes, including *Lophodermium conigenum* (Brunaud) Hilitzer, and *Rhizosphaera kalkhoffii* Bubák. *Lecanosticta acicola* was consistently identified from BSNB symptomatic trees from Alabama and Arkansas, Mississippi, Louisiana, and those single sites in Georgia and Texas that displayed BSNB symptoms. The pathogen was not detected from needlecast affected trees from Florida, South Carolina, or the remaining sites in Georgia and Texas. An unidentified member of the family Mycosphaerellaceae was identified in diseased needles from multiple states, and sequences that matched to *Fusarium sporotrichioides* Sherb. were detected in samples from South Carolina and Florida. Although identical ESVs matching this *Fusarium* were found at multiple sites, it must be noted that ITS is not an ideal barcode for the identification of individual *Fusarium* spp. and for this reason precise identification by metabarcoding is not possible. The endophytic *Lophodermium australe* Dearn. was found only among diseased needles from stands in several states, mostly from Arkansas, Georgia, and Texas. Another unidentified *Lophodermium* species was detected primarily in diseased needles in nearly all states surveyed. Multiple members of the Teratosphaeriaceae were also detected from diseased needles more often than in asymptomatic foliage. Certain fungal taxa were recorded as associates of most samples regardless of symptom presentation, including *Alternaria alternata* (Fr.) Keissl., *Septorioides pini*-*thunbergii* (S. Kaneko) Quaedvl., Verkley & Crous, and an unidentified *Pestalotiopsis* sp. A number of Mycosphaerellaceae were also detected among various symptom presentation types, including *Acrodontium*, *Geastrumia*, *Ramularia*, and *Zasmidium* spp., as well as *Australosphaerella nootherensis* (Crous et al. 2019; Videira et al. 2017).

**Table 4:**
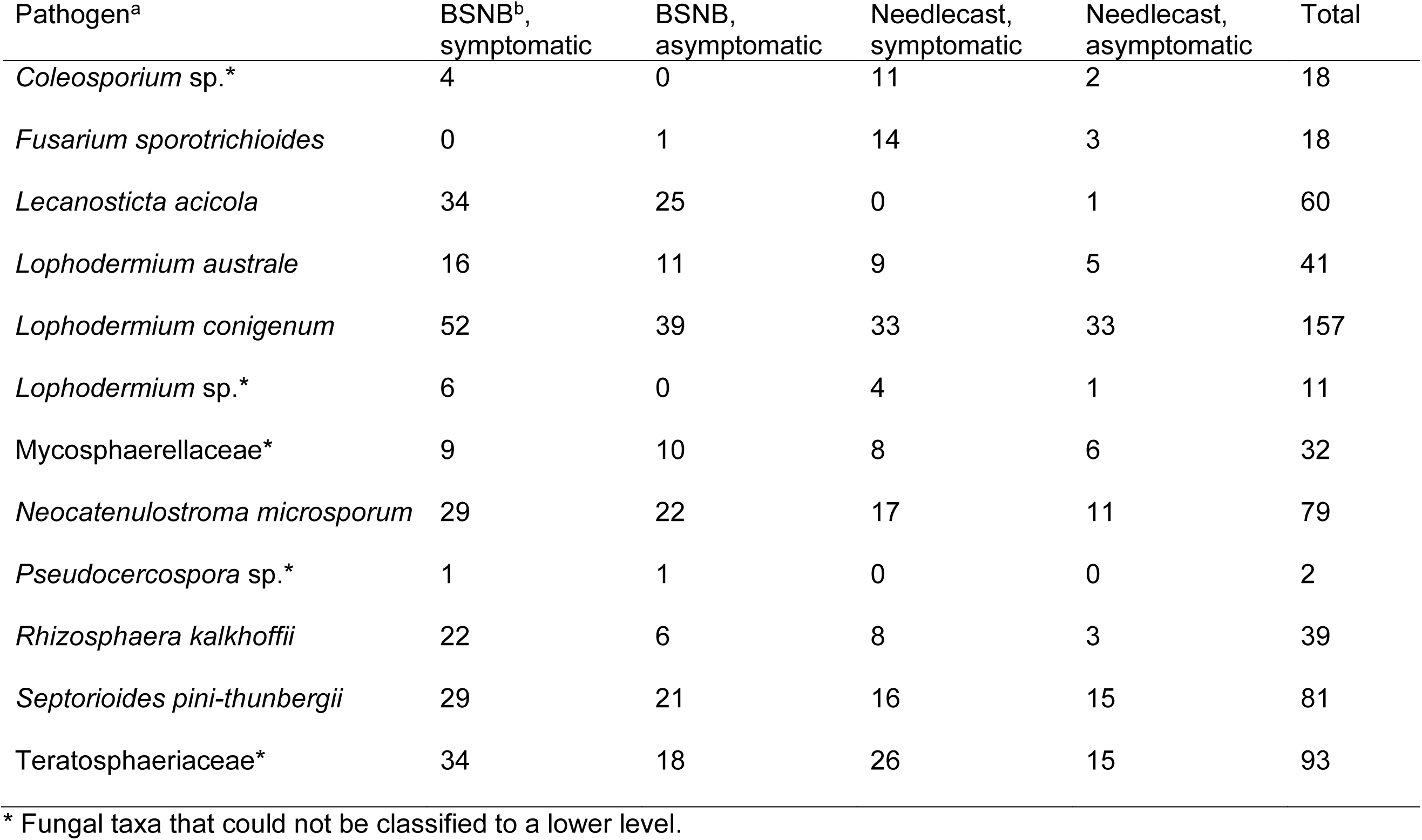
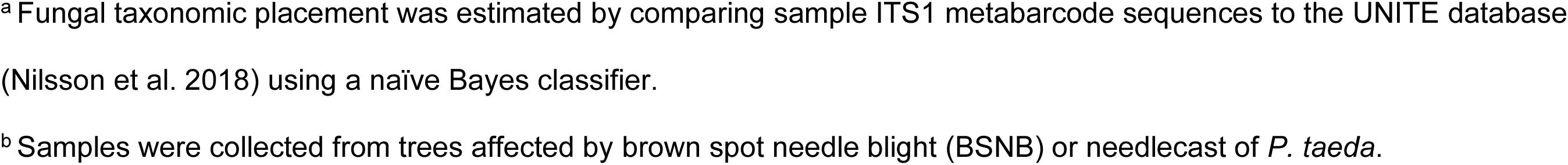
Fungi known to cause foliar disease in *Pinus* spp. identified in loblolly pine *(Pinus taeda*) needle samples that were symptomatic or asymptomatic, collected from trees that were affected by needle diseases.

| Pathogen<sup>a</sup> | BSNB<sup>b</sup>,<br>symptomatic | BSNB,<br>asymptomatic | Needlecast,<br>symptomatic | Needlecast,<br>asymptomatic | Total |
| --- | --- | --- | --- | --- | --- |
| *Coleosporium* sp.* | 4 | 0 | 11 | 2 | 18 |
| *Fusarium sporotrichioides* | 0 | 1 | 14 | 3 | 18 |
| *Lecanosticta acicola* | 34 | 25 | 0 | 1 | 60 |
| *Lophodermium australe* | 16 | 11 | 9 | 5 | 41 |
| *Lophodermium conigenum* | 52 | 39 | 33 | 33 | 157 |
| *Lophodermium* sp.* | 6 | 0 | 4 | 1 | 11 |
| Mycosphaerellaceae* | 9 | 10 | 8 | 6 | 32 |
| *Neocatenulostroma microsporum* | 29 | 22 | 17 | 11 | 79 |
| *Pseudocercospora* sp.* | 1 | 1 | 0 | 0 | 2 |
| *Rhizosphaera kalkhoffii* | 22 | 6 | 8 | 3 | 39 |
| *Septorioides pini-thunbergii* | 29 | 21 | 16 | 15 | 81 |
| Teratosphaeriaceae* | 34 | 18 | 26 | 15 | 93 |

\* Fungal taxa that could not be classified to a lower level.
<sup>a</sup> Fungal taxonomic placement was estimated by comparing sample ITS1 metabarcode sequences to the UNITE database (Nilsson et al. 2018) using a naïve Bayes classifier. <sup>b</sup> Samples were collected from trees affected by brown spot needle blight (BSNB) or needlecast of *P. taeda*.

### Fungal indicators of needle disease symptoms

Nine ESVs, corresponding to six taxonomic groups, were identified as indicators of BSNB or needlecast symptoms (Table 5). *Lecanosticta acicola* and *Neocatenulostroma microsporum* (Taylor & Crous) Quaedvl. & Crous were identified as strong predictors of the observed BSNB symptoms, while indicators of needlecast included multiple ESVs assigned to the *P. taeda* needle endophyte and weak pathogen *Lophodermium conigenum,* the cereal crop and pine pathogen *Fusarium sporotrichioides* (Ivanová et al. 2016), and unidentified members of the genus *Coleosporium* and the family Teratosphaeriaceae (Figure 3).

**Figure 3:**
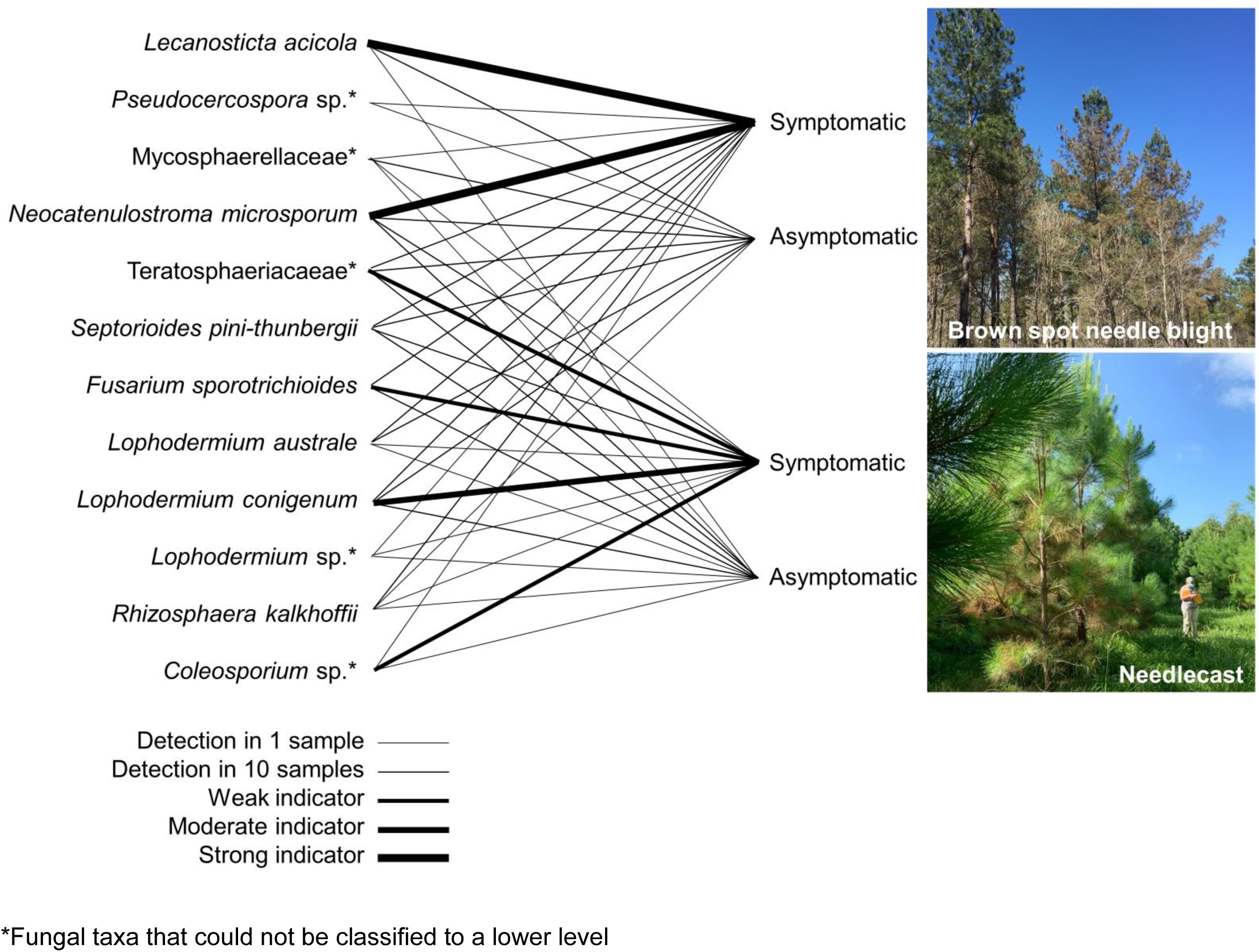
Fungi known to cause foliar disease in *Pinus* spp. identified in loblolly pine *(Pinus taeda*) needle samples that were symptomatic or asymptomatic, collected from trees that were affected by needle diseases brown spot needle blight or needlecast of *P. taeda*. Fungal taxonomic placement was estimated by comparing sample ITS1 metabarcode sequences to the UNITE database (Nilsson et al. 2018) using a naïve Bayes classifier. Line thickness is based on strength of association to needles with the observed symptom type from trees with brown-spot needle blight or needlecast based on presence and, for diseased samples, indicator value (IndVal). From thinnest to thickest, the lines represent taxa found in at least 1 sample, those found in at least 10 samples, weak indicators of the symptom type (IndVal > 0.50), moderate indicators (IndVal > 0.60) and strong indicators (IndVal >0.80).

**Table 5:**
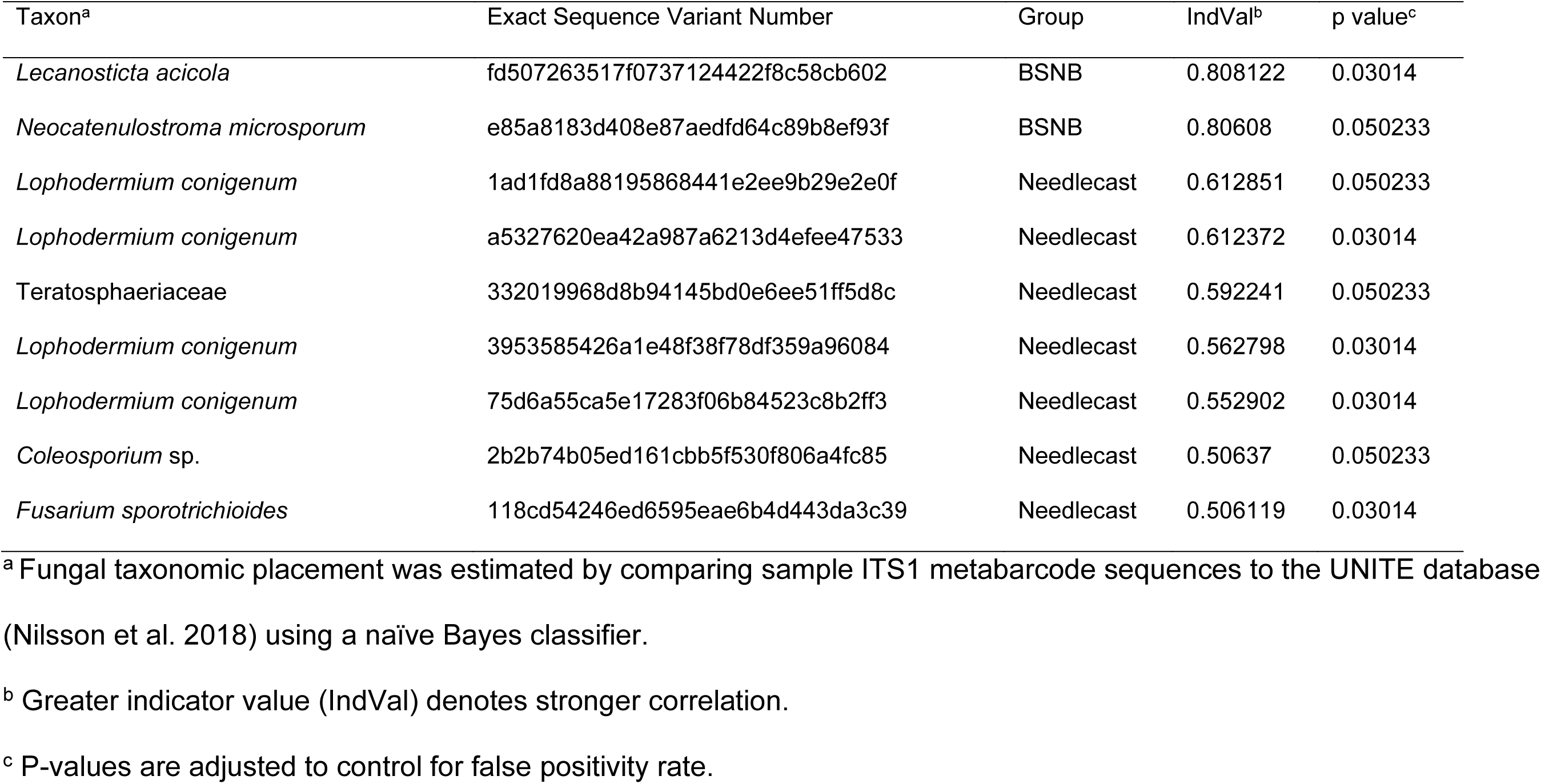
Indicator species of loblolly pine (*Pinus taeda*) needles displaying symptoms of brown spot needle blight (BSNB) and needlecast.

| Taxon<sup>a</sup> | Exact Sequence Variant Number | Group | IndVal<sup>b</sup> | p value<sup>c</sup> |
| --- | --- | --- | --- | --- |
| *Lecanosticta acicola* | fd507263517f0737124422f8c58cb602 | BSNB | 0.808122 | 0.03014 |
| *Neocatenulostroma microsporum* | e85a8183d408e87aedfd64c89b8ef93f | BSNB | 0.80608 | 0.050233 |
| *Lophodermium conigenum* | 1ad1fd8a88195868441e2ee9b29e2e0f | Needlecast | 0.612851 | 0.050233 |
| *Lophodermium conigenum* | a5327620ea42a987a6213d4efee47533 | Needlecast | 0.612372 | 0.03014 |
| Teratosphaeriaceae | 332019968d8b94145bd0e6ee51ff5d8c | Needlecast | 0.592241 | 0.050233 |
| *Lophodermium conigenum* | 3953585426a1e48f38f78df359a96084 | Needlecast | 0.562798 | 0.03014 |
| *Lophodermium conigenum* | 75d6a55ca5e17283f06b84523c8b2ff3 | Needlecast | 0.552902 | 0.03014 |
| *Coleosporium* sp. | 2b2b74b05ed161cbb5f530f806a4fc85 | Needlecast | 0.50637 | 0.050233 |
| *Fusarium sporotrichioides* | 118cd54246ed6595eae6b4d443da3c39 | Needlecast | 0.506119 | 0.03014 |

<sup>a</sup> Fungal taxonomic placement was estimated by comparing sample ITS1 metabarcoding sequences to the UNITE database (Nilsson et al. 2018) using a naïve Bayes classifier.
<sup>b</sup> Greater indicator value (IndVal) denotes stronger correlation.
<sup>c</sup> P-values are adjusted to control for false positivity rate.

### Fungal community composition in relation to geographic location and needle disease symptom presentation

The fungal community composition of pine needle samples differed by symptom presentation type (F = 2.6903, p < 0.001) and sampling location (F = 3.6284, p < 0.001), and an interaction effect between these factors was observed (F = 1.9348, p < 0.001) (Table 6). The PERMANOVA results indicate that the fungal community membership of *P. taeda* needles is distinct between needles with different disease symptoms and between different geographic locations.

**Table 6:** Results of a permutational multivariate analysis of variance to assess differences in fungal community composition between needle disease samples.

| Factor | DF<sup>a</sup> | SS<sup>b</sup> | R<sup>2</sup> | F | Pr (>F) |
| --- | --- | --- | --- | --- | --- |
| Symptom type | 3 | 2.808 | 0.04163 | 2.6903 | 0.001 |
| State | 7 | 8.836 | 0.13100 | 3.6284 | 0.001 |
| Symptom:State | 9 | 6.058 | 0.08981 | 1.9348 | 0.001 |
| Residual | 143 | 49.749 | 0.73756 |  |  |
| Total | 162 | 67.450 | 1.00000 |  |  |

<sup>a</sup> Degrees of freedom
<sup>b</sup> Sum of squares

A mean of 48.33 ESVs were present in each sample. Taxa richness was lower in asymptomatic needles from BSNB trees compared to symptomatic needles in both BSNB (p = 0.019) and needlecast-affected trees (p = 0.020) (Figure 4). Taxa richness did not vary among sampling locations (Figure 5).

**Figure 4:**
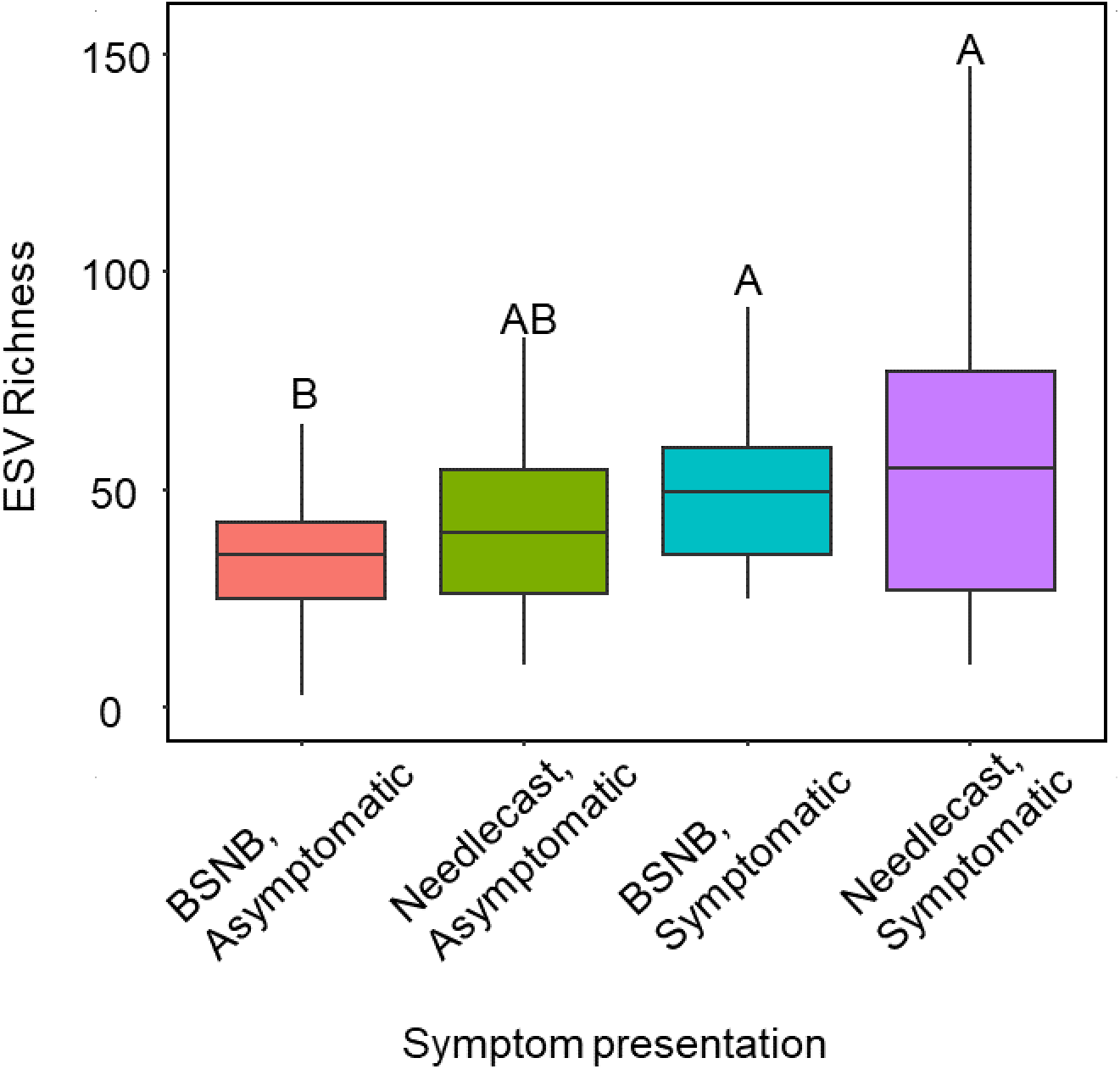
Taxa richness by pine needle symptom presentation type, as measured in number of exact sequence variants (ESVs) identified from ITS1 metabarcode data. Each box indicates 25th-75th percentile of ESVs by symptom type, and the ends of whiskers indicate 5th and 95th percentile. The center line represents the mean richness value. Letters indicate symptom types that differed significantly.

**Figure 5:**
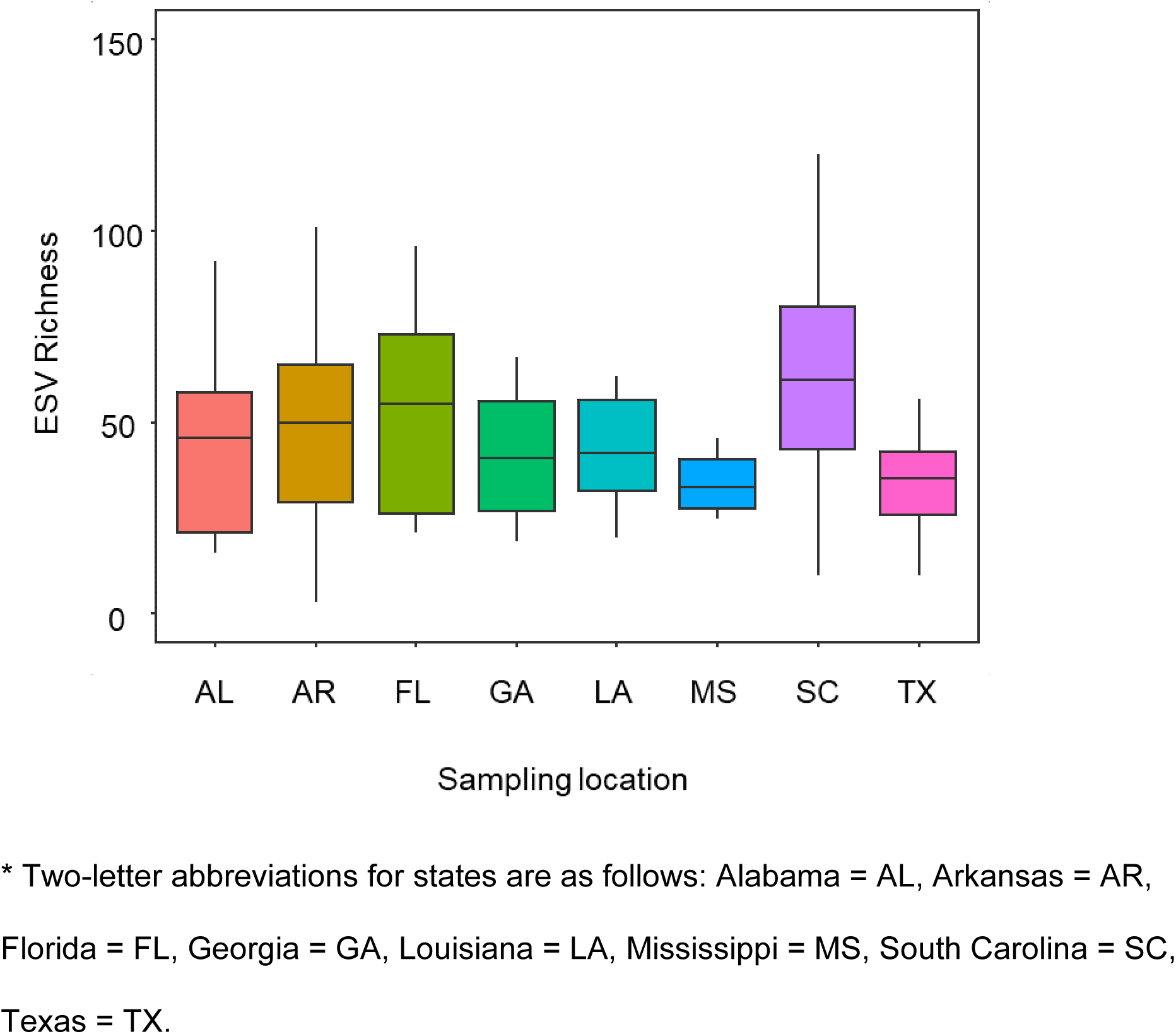
Taxa richness by geographic sampling location*, as measured in number of exact sequence variants (ESVs) identified from ITS1 metabarcode data. Each box indicates 25th-75th percentile of ESVs by sampling location, and the ends of whiskers indicate 5th and 95th percentile. The center line represents the mean richness value. No significant differences were detected among states.

## Discussion

In this study, we analyzed needles from *P. taeda* affected by foliar diseases to identify fungal pathogens associated with outbreaks in the southeast United States. We hypothesized that the BSNB pathogen *L. acicola* would be present among outbreaks, that certain pathogenic fungi would be identified as indicators of disease, and that fungal communities of needles collected from outbreak sites would differ between symptom presentation types and sampling locations. Through this investigation, we identified two specific and diverse foliar diseases impacting *P. taeda* in the southeast US, BSNB and needlecast. Our analysis of the fungal communities confirmed the presence of foliar pathogens, identified the different fungal indicators for each disease, and revealed that disease is linked to differences in community composition and diversity.

This study confirms the presence of *L. acicola* in samples from impacted stands in Alabama and Arkansas, supporting previous reports of the pathogen in association with BSNB damage and further indicating that *L. acicola* is a growing threat to *P. taeda* forests (Datta et al. 2021; Meinecke et al. 2021). However, the brown spot pathogen was not detected in needlecast symptomatic samples from Georgia, South Carolina, or Florida. While *L. acicola* is reported in association with diseased *P. taeda* all throughout the southeast US, most observations are recorded in heavily infested sites in the Gulf states of Alabama, Mississippi, and Louisiana, where the pathogen is reported among major BSNB outbreaks. The sporadic presence of *L. acicola* from eastern sites, coupled with different symptom presentation, suggests that other pathogens are driving the needlecast outbreaks in these states.

The species-specific LAMP assay was able to identify *L. acicola* infections at most BSNB symptomatic sites where the fungus was confirmed to be present using DNA metabarcoding, although it detected the pathogen in a lower proportion of symptomatic samples (41.7%) than metabarcoding, and rarely in asymptomatic needles. The LAMP-based approach was selected as an inexpensive and rapid screening tool. However, it has a limit of detection comparable to end-point PCR at 0.128 pg/µL (Aglietti et al. 2021) and was thus expected to achieve a lesser degree of sensitivity compared to the Illumina amplicon-based metabarcoding, whose sensitivity is comparable to that of nested PCR reactions due to the two rounds of PCR used to prepare libraries. The results of this study confirm that LAMP can serve as a sustainable option for routine diagnostic screenings when a molecular level of confirmation is needed in a short amount of time, with the caveat that multiple samples should be tested to compensate for its limited sensitivity.

The ISAs identified several other taxa as indicators of either BSNB or needlecast symptom presentation. While the precise taxonomic identities of each ESV must be considered putative, a limited set of ESVs were significantly associated with the different disease patterns. A single ESV matching to *Neocatenulostroma microsporum*, for instance, was a strong indicator of BSNB symptoms. While the pathogenicity of *N. microsporum* on *P. taeda* is not known, the closely related *N. germanicum* (Crous & U. Braun) Quaedvl. & Crous is a causal agent of needle blight on *Pinus* spp. in east Europe and is known to co-occur with *L. acicola* and *Dothistroma* spp. in symptomatic tissues (Markovskaja et al. 2016). Experiments must be conducted to determine whether *N. microsporum* may be a pathogen on *P. taeda* or if co-infection with *L. acicola* may yield synergistic or antagonistic interactions. *Fusarium sporotrichioides,* which we identified as a moderate predictor of needlecast symptoms is a well-known mycotoxigenic fungus and cereal crop pathogen and was recently reported in association with wilt and defoliation on *P. ponderosa* in Slovakia (Ivanová et al. 2016). As *Fusarium* spp. have not been reported to attack *P. taeda* foliage, future assessments of the role of *F. sporotrichioides* in needlecast symptom development on this host are needed.

Multiple ESVs that were assigned to *Lo. conigenum* were moderate indicators of needlecast. It must be noted that the endophyte *Lo. conigenum* cannot be differentiated from the more aggressive *Lo. pinastri* (Shrad.:Fr.) Chev. based on ITS sequence alone. Typically regarded as opportunistic weak pathogens in the region, *Lo. conigenum* and *L. pinastri* are commonly observed and associated with damage to southern pines, although serious damage is rare (Barnard and Ash 1998; Oono et al. 2015).

Sequences that were classified as *Lo. conigenum* made up the majority of reads in most samples. The greater number of ESVs identified relative to the other taxa of interest likely speaks to the sequence diversity that was visible due to the greater depth of reads assigned to *Lo. conigenum*. However, the identification of select ESVs as predictors of disease may indicate signatures of virulence among those taxa, the presence of cryptic species diversity, or intraspecific variation in the ITS1 subregion. Lastly, unidentified members of the family Teratosphaeriaceae and the genus *Coleosporium* were significantly associated with needlecast. Unidentified Teratosphaeriaceae are known associates of spring needlecast on *Pinus* spp. in Tasmania, along with *Lo. pinastri* (Prihatini et al. 2015). *Coleosporium* spp. are the causal agents of needle rusts on *Pinus* spp (Aime et al. 2017). Rust damage was not apparent on needle samples, and although rust is not likely to be an outright cause of the observed necrotic damage, the possibility of interactions among multiple pathogens remains. Additional work is needed to understand how these taxa contribute to needlecast outbreaks on *P. taeda*. The multitude of indicator taxa suggests that the emerging *P. taeda* needle diseases are complex, and while there are likely major pathogens responsible for most of the damage, there are possibly interactions between multiple organisms that are not yet understood.

Many common opportunistic pathogens that are known associates of *P. taeda*, as well as several unexpected taxa, were identified among those detected in needle samples. In addition to those highlighted in the ISAs, the metabarcoding analysis detected *R. kalkhoffii, S. pini-thunbergii,* and unidentified *Lophodermium* and *Pseudocercospora* spp. As with *Lophodermium* spp., *R. kalkhoffii* is a weak pine pathogen associated with brief, sporadic, and episodic disease events on southern pines (Barnard & Ash, 1998). *Septorioides* spp. are known needle pathogens and endophytes of *Pinus* (Quaedvlieg et al. 2013). Notably, *S. pini-thunbergii* is an associate of needle blight on Japanese black pine (*P. Thunbergii* Parl.) (Kaneko et al. 1989) and an endophyte of *P. densiflora* Siebold & Zucc. (Yoo and Eom 2012), while another species in the genus, *S. strobi* Wyka & Broders, is frequently identified in cases of WPND (Wyka and Broders 2016). The detection of a *Pseudocercospora* sp. requires additional investigation, as *P. pini-densifloriae* (Hori & Nambu) Deighton is the causal agent of brown needle blight on *Pinus* and a quarantine pest in many countries (EFSA et al. 2017). Other members of the genus *Pseudocercospora* cause leaf spots on agricultural crops (Ávila et al. 2019; Crous et al. 2021). Finally, other members of the family Mycosphaerellaceae, including *Acrodontium*, *Geastrumia*, *Ramularia*, and *Zasmidium* spp, and the *Corymbia intermedia* (Baker) Hill & Johnson pathogen *Australosphaerella nootherensis* were found across different locations and symptom types. An unidentified species in the Mycosphaerellaceae was also identified in needlecast symptomatic samples from Georgia. The family Mycosphaerellaceae includes many of the most important foliar pathogens, including *Lecanosticta, Dothistroma*, and *Mycosphaerella* spp. (Crous et al. 2019; Videira et al. 2017). While these taxa were not widely detected in this study or identified as indicators of needle disease symptoms, their mere presence among needle disease outbreaks is alarming and warrants further investigation.

Interestingly, *L. acicola* was not detected by LAMP nor by metabarcoding in several samples that displayed BSNB symptoms. This is possibly due to the low proportion of *L. acicola* DNA relative to host DNA in the raw DNA extracts, possible primer bias, or to data loss during processing. After quality filtering and random subsampling, *L. acicola* reads comprised a low proportion of total reads in infected samples (0.06% to 14.3%), bordering on the lower limit of detection. Measures to ensure even sampling are known to reduce the sensitivity of NGS-based detection (Tedersoo *et al*. 2022). However, these methods enable high-resolution surveys of fungal communities and the detection of fastidious fungi that would be otherwise impossible using traditional culture-based techniques.

The needle samples analyzed in this study significantly differed in their fungal community composition by both symptom type and by sampling location. Foliar diseases can alter endophyte community assemblies, as recently shown in cases of *Lophodermella* infection (Ata et al. 2022), and differences in community composition by location are likewise unsurprising considering the vast geographic variation among study sites. Interestingly, ESV richness was greater in BSNB and needlecast symptomatic needles when compared to the BSNB asymptomatic needles but did not differ between any other pairs of symptom types or by location. This is likely due to opportunistic pathogens and saprophytes invading weakened and dead tissues.

The ISAs demonstrate a strong correlation between *L. acicola* presence and BSNB symptoms. These findings build upon the growing body of work that demonstrates a clear association between *L. acicola* infection and the damaging outbreaks observed in the western portions of the *P. taeda* range (Datta et al. 2021; Meinecke et al. 2021). *Lecanosticta acicola* was detected in the majority of BSNB symptomatic needles (71.4%) and was consistently detected among asymptomatic needles from the same hosts (60.0%). It is not surprising that *L. acicola* was identified in asymptomatic needles from BSNB affected trees, as the fungus may colonize without eliciting disease for some time (van der Nest et al. 2019b) and latent infections are likely common in active outbreak sites.

The simultaneous eruption of separate needle diseases with different associated fungi speaks to a broader pattern of increasing foliar pathogen risk. This pattern is possibly due to climactic shifts that create conditions favorable to disease development, those that increase host tree stress, or both (Pandit et al. 2020). In fact, the potential distribution of *L. acicola* is expected to expand to include over half of all global pine cover due to the warming climate (Ogris et al. 2023). This study identified several likely pathogens as disease predictors in heavily damaged stands, including *L. acicola,* which is a species of growing concern worldwide (Tubby et al. 2023; van der Nest et al. 2019b). On its own, our data is insufficient to propose causal relationships between the observed large-scale damage and fungi detected in needle samples, as this relationship must be proven through controlled pathogenicity tests and the fungal identities are putative (Fredericks and Relman 1996; Severns and Sykes 2020). However, this study provides a robust and condensed list of organisms that should be considered in future studies on causal agents of the unprecedented *P. taeda* needle disease outbreaks in the southeast United States. Additional work is also needed to design broadly applicable molecular diagnostic approaches that maximize the generality and high-throughput capability of next generation sequencing-based metabarcoding with the precision of classical fungal barcoding. Human and plant disease diagnostics studies are currently developing methods that utilize the enrichment of single or multiple barcode loci in tandem with long read sequencing (Zhang et al. 2020; Yu et al. 2023) and offer the potential to greatly enhance pathogen identification speed and precision. In conclusion, this study is the first comprehensive work to identify potential pathogens associated with *P. taeda* needle disease across the southeast United States. By leveraging whole fungal community profiles, we identified fungi that predict disease presentation, facilitating the future confirmation of causal agents. Precise identification of the pathogens involved will be critical to the future development of strategies to manage disease whether by the use of fungicides or biocontrol applications, or through silvicultural intervention or resistance breeding.

